# Deformed Wing Virus spillover from honey bees to bumble bees: a reverse genetic study

**DOI:** 10.1101/2019.12.18.880559

**Authors:** Olesya N Gusachenko, Luke Woodford, Katharin Balbirnie-Cumming, Ryabov Eugene V Ryabov, David J Evans

**Affiliations:** Evans laboratory, Biomedical Sciences Research Complex, University of St. Andrews, North Haugh, St. Andrews, UK; USDA-ARS Bee Research Laboratory, Beltsville Agricultural Research Center, Beltsville, MD 20705, USA

## Abstract

Deformed wing virus (DWV) is a persistent pathogen of European honey bees and the major contributor to overwintering colony losses. The prevalence of DWV in honey bees has led to significant concerns about spillover of the virus to other pollinating species. Bumble bees are both a major group of wild and commercially-reared pollinators. Several studies have reported pathogen spillover of DWV from honey bees to bumble bees, but evidence of a true sustained viral infection has yet to be demonstrated. Here we investigate the infectivity and transmission of DWV in bumble bees using the buff-tailed bumble bee *Bombus terrestris* as a model. We apply a reverse genetics approach combined with controlled laboratory conditions to detect and monitor DWV infection. A novel reverse genetics system for three representative DWV variants, including the two master variants of DWV - type A and B - was used. Our results directly confirm DWV replication in bumble bees but also demonstrate striking resistance to infection by certain routes. Bumble bees may support DWV replication but it is not clear how infection could occur under natural environmental conditions.

## Introduction

Deformed wing virus (DWV) is a widely established pathogen of the European honey bee, *Apis mellifera*. In synergistic action with its vector — the parasitic mite *Varroa destructor* — it has had a devastating impact on the health of honey bee colonies globally. As the primary managed insect pollinator, honey bees are of high ecological and economic value and contribute an estimated 30-50% of mobile pollination activity [1, 2]. Two-thirds of all colonies in the USA (∼1.6 million hives) are transported to California in February/March for almond pollination [3]. Inevitably, transporting bees also transports their pathogens. This, coupled with the local pathogen density associated with ∼50 000 bees in a single hive, has raised concerns about pathogen spillover from managed honey bees to other pollinators [4]. DWV was found as a frequent component of pollen pellets [5] and is also present in bee faeces [6], suggesting honey bee foragers and colonies could facilitate horizontal virus transmission to the wider pollinator community. Significantly, DWV RNA has been detected in many insects sharing the environment with managed honey bees, including Asian bee species and wild bees, cockroaches, ants, wasps, and bumble bees [4, 5, 7–19]. Due to their extended activity at lower temperatures (compared to honey bees) bumble bees are considered particularly important pollinators in temperate and subarctic climates [20, 21] and are the only other insect managed for commercial-scale pollination [21]. As a consequence, the potentially negative impact of extensive honey bee management and failing pathogen control on co-located *Bombus* species has received significant attention.

Following a report that DWV was detected in symptomatic *Bombus terrestris* and *Bombus pascuorum* individuals with deformed wings [7] there have been several studies of DWV prevalence in a wide range of *Bombus* species [4, 5, 11, 12, 22–24]. In *Varroa*-infested honey bee colonies DWV levels can exceed 10^11^ genome copies per bee [25], with considerable potential for environmental contamination. DWV-positive *Bombus sp*. have been shown to correlate to areas with high DWV prevalence in *Apis* [4, 23, 24]. The majority of screening studies have used RT-PCR for DWV detecting both positive and negative strand DWV RNA in environmental bumble bee samples (reviewed in [26]). However, the near ubiquitous presence of managed hives, the honey bee – and consequently pathogen – density around hives, and the foraging range of *Apis* mean that DWV is likely widespread [5]. Although detailed analysis of DWV replication requires detection of the negative strand intermediate of replication it should be noted that this alone is not formal proof that the virus is replicating. There remains the possibility that negative strands are present as carry-over in *Apis*-derived cellular material ingested by bumble bees.

The name DWV is currently attributed to an evolving complex of closely related viruses, which includes DWV-A [27], Kakugo virus [28], Varroa destructor virus-1 (VDV-1; also referred to as DWV-B [29–31]) and a range of recombinants between DWV-A and -B [32–35]. All viruses exhibit at least 84% identity at the nucleotide level and 95% identity at the protein level [32, 35, 36]. The sensitivity of current diagnostic methods means DWV detection in environmental samples is regularly reported, with different DWV variants identified in *Bombus* [4]. Far less frequent are studies investigating potential routes of transmission from *Apis* to *Bombus*, or the subsequent replication of DWV in bumble bees. Due to the absence of a suitable cell line for *in vitro* propagation, laboratory-based assays have been limited to the application of field-sourced virus. Infectivity of DWV obtained from field honey bee samples was tested via inoculations of adult *Bombus terrestris* [4, 22]. It was reported that a DWV complex containing both DWV-A and -B is infectious when fed at high concentrations - 10^9^ GE of virus per bee [4].

We have used a reverse genetic (RG) approach to generate near-clonal populations of genetically tagged DWV-A, -B and a B/A hybrid after transfection of honey bees with *in vitro* transcribed RNA. A similar system has recently been reported for DWV-A [37–39]. Using RG-derived DWV inocula we address the question of DWV pathogenesis and likely transmission routes in *Bombus terrestris* at both the individual and colony level. Using this strategy we provide direct evidence of DWV replication in this pollinator via virus feeding and injection. Importantly, adult *Bombus terrestris* appear resistant to productive infection by DWV orally and do not exhibit the developmental defects characteristic of DWV infection and replication in honey bees.

## Results

### Infectivity of DWV RNA and virus in injected honey bee and bumble bee pupae

In order to test infectivity of DWV variants in controlled laboratory experiments we have developed RG systems for DWV-A and -B master variants. DWV cDNAs were built using gene synthesis based upon published sequences [27, 40, 41] and that of a recombinant DWV reported earlier [25]. The RG constructs design is similar to a recently reported system for DWV-A [37] with a full-length DWV cDNA placed under transcriptional control of a T7 promoter. RNA generated *in vitro* is directly injected into honey bee pupae from which infectious virus is recovered. To allow discrimination from any endogenous DWV we engineered synonymous mutations creating new restriction sites, allowing the unambiguous identification of the RG genomes. The following nomenclature was used for the generated viruses: “VDD” - type A DWV, “VVV” - type B, “VVD” - B/A recombinant (Figure 1A).

**Figure 1.**
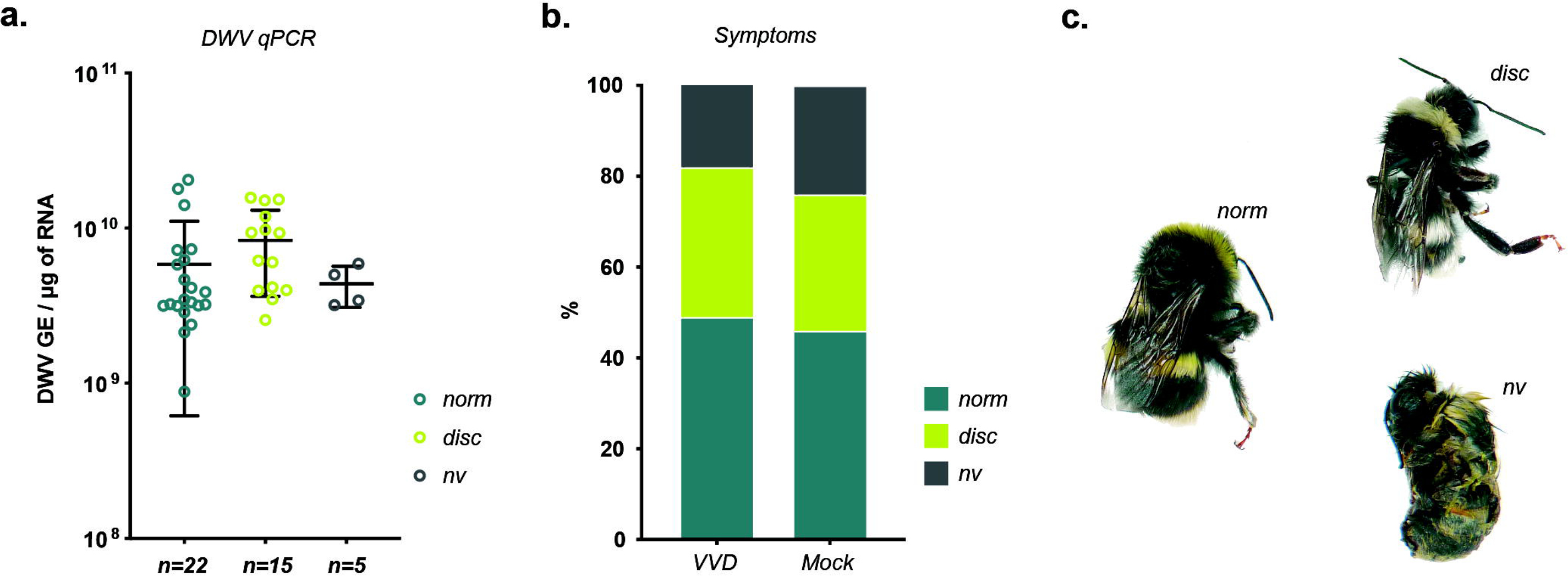
Reverse genetics (RG) system for three DWV variants. **A**. Diagram showing identical regions between genomic RNA sequences of master DWV variants (DWV-A - GenBank AJ489744 and DWV-B - GenBank AY251269.2) and three RG clones (VVV, VDD and VVD) derived from the cloned cDNA of the recombinant DWV - VVD (GenBank HM067438.1); identical sequence regions are shown in black, restriction sites tags introduced into cDNA of each variant are indicated in italics, DWV genomic RNA organization is presented below to help interpretation. **B**. Detection of RG DWV RNA by RT-PCR in honey bee pupae injected with *in vitro* synthesized RNA transcripts: “VDD”, “VVV”, and “VVD” - PCR products from pupae injected with corresponding full-length RNA, “VVDtr” - PCR from pupae, injected with truncated VVD RNA, “+” and “-” - PCR controls, “M” - molecular weight DNA marker; restriction digest - verification of the RG origin of detected DWV by the digest of the PCR products at artificially introduced restriction sites. **C**. Detection of RG DWV in bumble bee pupae injected with VVD RNA and with virus stock obtained from RNA-injected pupae - “VVDvir”; “VVD RNAtr” - RT-PCR from pupae injected with truncated RNA, “Mock” - PBS-injected pupae, “+” - positive PCR control for DWV, “M” - molecular weight DNA marker; RG origin of the PCR products for all DWV-positive samples was confirmed by digest with *HpaI* restriction enzyme, amplification of the actin mRNA product was used as an indicator of RNA integrity and loading control.

In temperate climates honey bee brood is only available for ∼50% of the year and *in vivo* research is of necessity seasonal. Although the DWV IRES retains partial activity in at least one cell line of non-honey bee origin (*Lymanthria dispar* LD652Y cells) [42], attempted infection of those cells with DWV or transfection of *in vitro-*generated DWV RNA did not result in virus replication (OG, unpublished data). This impediment to DWV studies prompted us to investigate the recovery of clonal stocks of DWV in bumble bees (*Bombus terrestris audax*) injected with viral RNA. Commercially farmed bumble bee colonies are available year-round and, in our preliminary studies, are free of DWV RNA (data not shown). Four of six RNA-injected bumble bee pupae tested positive for DWV, while all honey bee pupae injected with full-length RNA were shown to be DWV-positive (Figure 1B, C). The RG origin of DWV in injected samples was confirmed by restriction digest of PCR products using endonucleases specific for the introduced genetic tag (Figure 1B,C - restriction digest). The remaining tissue from RNA-injected samples was used to prepare crude DWV stocks. RT-qPCR analysis showed that the *Apis-*derived stocks contained between 10^7^-10^9^ genome equivalents (GE) of DWV/μl, while *Bombus*-derived inoculum contained 3.75×10^5^ GE/μl. For comparison, DWV levels in mock-injected *Apis* pupae did not exceed 10^5^ GE DWV/μg of RNA (representing the endogenous viral load) and *Bombus* pupae had no detectable DWV sequences in mock-inoculated samples.

DWV extracted from *Bombus* was infectious when re-inoculated by injection to *Bombus* pupae. We observed no differences in the infectivity of *Apis-* or *Bombus-*derived DWV (Figure S1) and, at the genome level, no obvious signs of adaptation following comparison of the parental cDNA sequence with NGS data from the second passage of *Bombus*-derived DWV (Figure S2).

In honey bees, both DWV-A and -B are pathogenic when inoculated, though recent studies have produced contradictory results when comparing their relative virulence [29, 41, 43]. We therefore tested infectivity of DWV types A, B and a B/A hybrid in bumble bee brood and adults. As our primary interest was to investigate the potential for DWV spillover from infected honey bees we used DWV inoculates prepared from RNA-injected honey bee pupae for all further experiments.

White-eyed (1 day post molting) or purple-eyed (5.5 days) bumble bee pupae were injected with 10^3^ or 10^6^ GE of each DWV variant, and virus levels quantified 48 hours post-inoculation. In all cases we observed a 10^2^-10^4^ increase in total DWV load compared to the inoculated amount, providing clear evidence for replication (Figure 2A). More DWV accumulated in older pupae but this was only statistically significant for 10^6^ GE of the VVV variant (Tukey’s multiple comparisons test, P=0.03).

**Figure 2.**
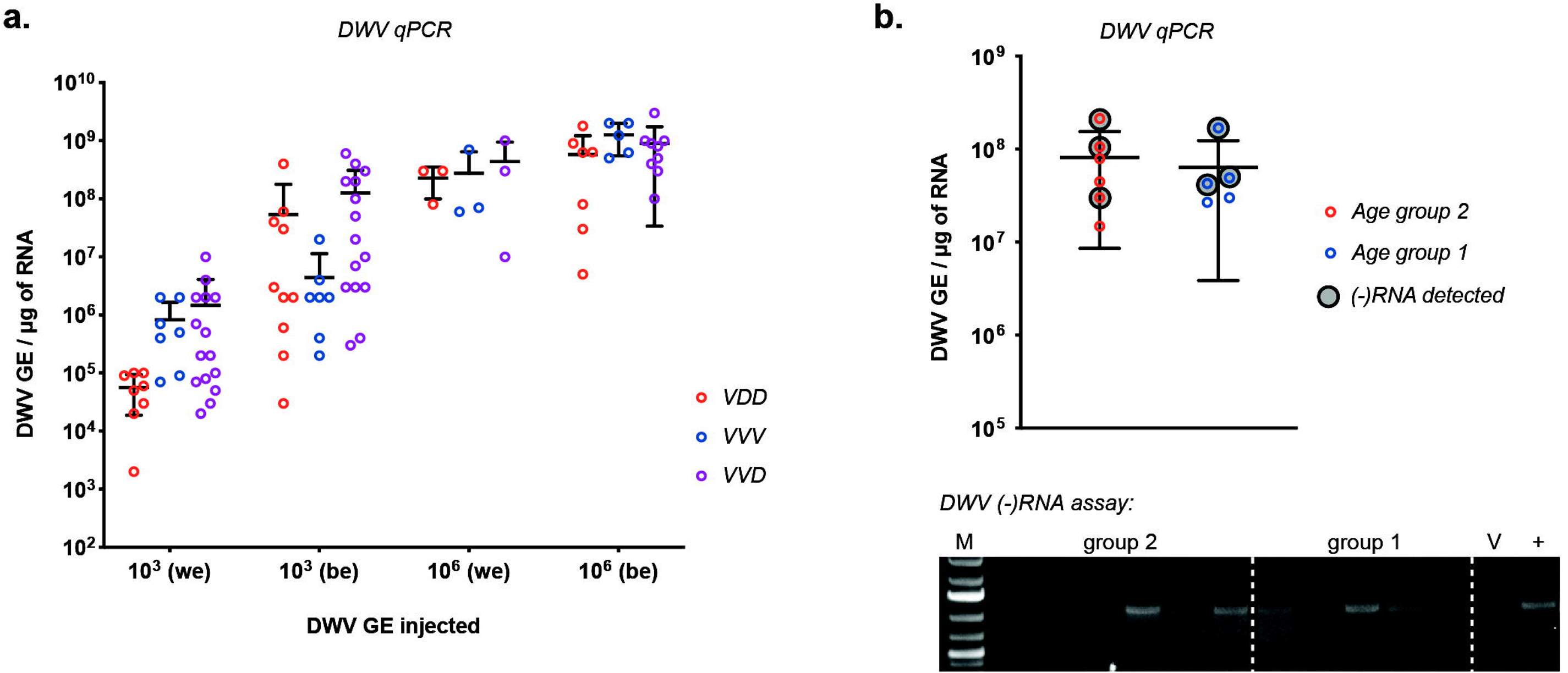
Inoculation of bumble bee pupae and larvae with RG-DWV. **A**. RT-qPCR analysis of DWV accumulation in bumble bee pupae injected with VVV, VDD and VVD DWV. Pupae were injected at white-eyed - “we” - and purple-eyed - “pe” - stages and analyzed 48 h post injection. Each value corresponds to an individual sample analyzed, black lines show mean ±SD. **B**. Detection of DWV RNA in bumble bee larvae from two different age groups fed with 10^8^ GE VVD DWV: qPCR analysis of DWV levels in individual larvae samples, black-circled values correspond to samples which produced a positive result in (-)RNA assay; each value corresponds to an individual sample analyzed, error bars show mean ±SD. DWV (-)RNA assay - strand-specific RT-PCR products run in 1% agarose gel, “+” - positive PCR control for DWV, “M” - molecular weight DNA marker, V - PCR from the DWV inoculum used for larvae feeding.

### *Bombus* larvae can be infected with DWV *per os*

Since bumble bee pupae are not parasitised by *Varroa* naturally-infected pupae must acquire the virus by prior exposure of larvae. We therefore investigated virus infection after feeding DWV to 1st and 2nd instar *Bombus* larvae (age 1 and 2 on Figure 2B respectively). Each larva individually received a single dose of diet containing 10^8^ GE of DWV on the first day of the experiment. DWV was detectable in all fed larvae on the 5^th^ day post inoculation, although virus levels were comparable to the amount administered in the diet. When analysed ∼50% of fed larvae had detectable levels of DWV negative strand RNA, absent from the inocula and indicative of virus replication.

### Investigation of colony scale transmission of DWV and adult infection *per os*

In honey bees, horizontal transmission of DWV occurs *per os* in larvae or adults, or when *Varroa* feed on pupae and adults. Whilst we show here that DWV can replicate in *Bombus sp*. pupae after direct injection, and in virus fed larvae, the route by which adult bumble bees could become infected remains unclear. We reasoned that two likely routes would be via direct oral exposure of adult *Bombus* to virus in the environment or indirectly following larval infection with virus carried by adult bees.

Groups of adult bumble bees received 10^7^ or 10^8^ DWV GE per bee via feeding with virus-supplemented sucrose solution. All control group bees remained viable during the experiment, and only one dead bee was found in each of the virus fed groups. Bees were analyzed 7 days after DWV feeding when none of the virus-fed samples tested positive for DWV (Figure 3). According to qPCR analysis of DWV-fed bees no sample contained greater than 10^5^ GE DWV per 1 μg of total RNA. Average RNA yield per bee did not exceed 50 μg and therefore the level of virus detected was 100-200-fold lower than the amount received. Hence, despite evidence that bumble bee larvae displayed susceptibility to DWV infection *per os*, feeding of adult bumble bees up to 10^8^ GE DWV per bee did not lead to infection or detectable replication.

**Figure 3.**
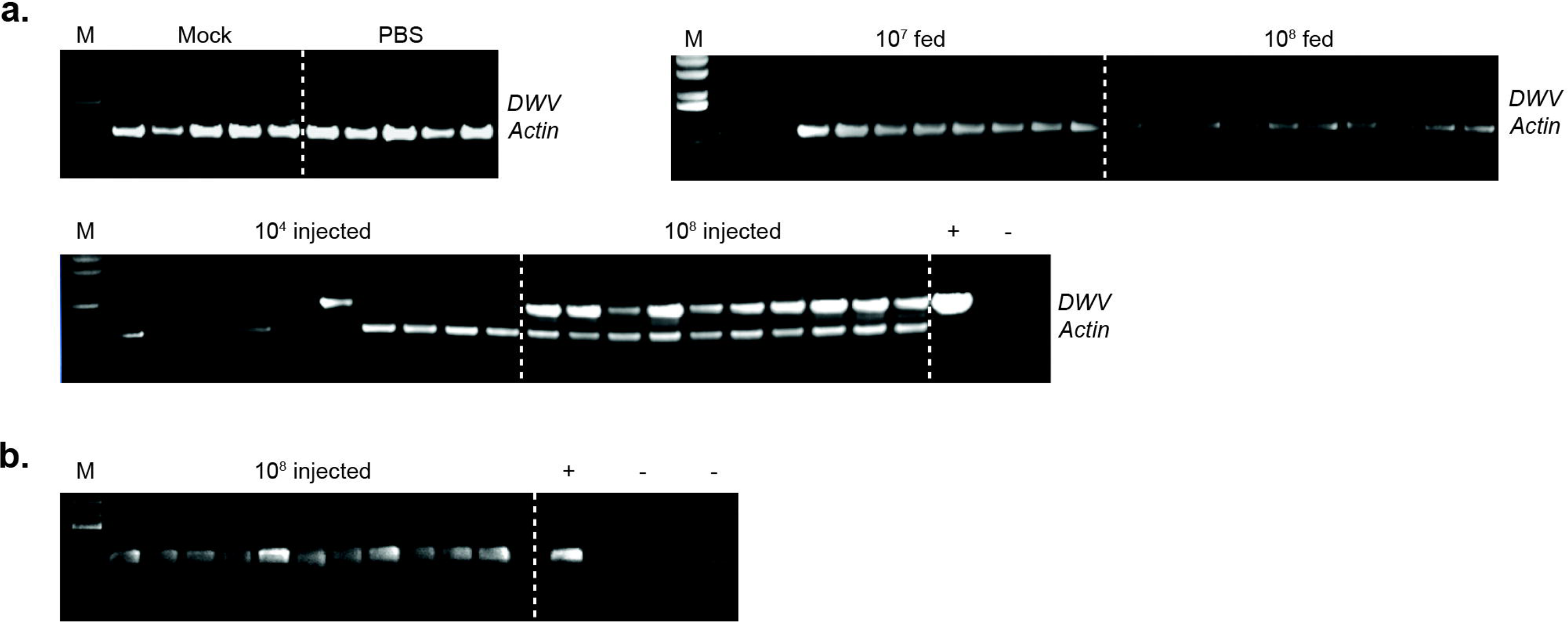
Detection of DWV in adult bumble bees. **A**. Detection of DWV RNA by RT-PCR in adult bumble bees after inoculation with VVD DWV via feeding or injections (10 samples from each group shown): “Mock” – non-treated bumble bees from the same colony, “PBS” - PBS-injected group (no virus), “10^7^ and 10^8^ fed” - bumble bees fed with sucrose solution containing 10^7^ or 10^8^ DWV GE of DWV per bee, “10^4^ and 10^8^ injected” - bumble bees injected with 10^4^ or 10^8^ DWV GE of DWV respectively. **B**. Detection of the DWV (-)RNA strand in adult bumble bees injected with 10^8^ DWV GE. “+” and “-” - positive and negative PCR controls, “M” - molecular weight DNA marker. Detection of the bumble bee actin RNA was used to assay the quality of extracted RNA. RG origin of the PCR products for all DWV-positive samples was confirmed by restriction enzyme digest.

We extended this study to investigate whole-nest inoculation with DWV. Three individual bumble bee nests were fed for 4-6 weeks with a sucrose solution supplemented with 2×10^8^ GE/adult bee/day of VVD, VVV or VDD DWV. All brood (pooled egg samples, 30 larvae, 47 pupae) and 30 adults from virus-exposed nests were screened for DWV using end-point RT-PCR assay and showed no positive results (data not shown).

### Direct inoculation of adult bumble bees

With no evidence that adult bumble bees could be infected when fed a DWV-supplemented diet, or that they were able to transfer virus to larvae, we performed direct virus injections of adult bumble bees to determine whether adults could support DWV replication. Two groups of adult bumble bees were intra-abdominally injected with 10^4^ or 10^8^ DWV GE per bee. Envisaging a possible impact of injection on viability, an additional group of bumble bees was injected with PBS solution only. 20-30% of injected bees died before termination of the experiment in each group. The remaining bumble bees were screened for DWV RNA 8 days post-inoculation. End-point PCR analysis indicated that 40% and 100% were DWV positive in the 10^4^-injected and 10^8^-injected groups respectively (Figure 3A), with qPCR analysis showing that the virus levels in these groups ranged between 2.74×10^4^-1.39×10^7^ and 4.82×10^6^-2.13×10^8^ GE/μg of total RNA respectively. Accumulation of DWV (-)RNA in DWV-positive samples was confirmed by strand-specific RT-PCR assay (Figure 3B). This demonstrates that DWV can replicate in adult *Bombus terrestris* after direct injection at 10^8^ GE per bee. We therefore proceeded to investigate if there were any pathogenic consequences of virus infection and replication.

### Pathogenicity of DWV in *Bombus terrestris*

DWV produces characteristic pathogenicity in honey bees and similar symptoms have been reported in bumble bees [7]. We injected 43 white-eyed bumble bee pupae with 10^6^ GE of DWV and maintained them through development in an incubator. In parallel, a group of 37 similarly-aged pupae were injected with PBS. All fully-developed bumble bees could be classified into one of three groups: normal phenotype, discolored with normal wings and non-viables, which did not eclose (Figure 4). Analysis by qPCR demonstrated that all DWV-inoculated bumble bees contained high levels of virus (8.8×10^8^-2.2×10^10^ GE/μg of RNA) with no significant differences in viral load between the three phenotypic groups (ANOVA, P=0.21) (Figure 4). DWV levels were 1-2 log_10_ greater than in pupae analysed 48 hours post-inoculation (Figure 2A) reflecting the additional time the virus had to replicate. This further supports the conclusion that pupae can be infected with DWV by direct inoculation. Strikingly, none of the eclosed bumble bees showed any signs of wing deformities that are characteristic of DWV infection of honey bees. Inspection of PBS-injected pupae showed that they could be separated into the same phenotypical groups; normal, discoloured and dead. There was no statistical difference between the proportions in each group of virus- or PBS-injected bumble bees (ANOVA, P>0.999). Upon analysis, none of the mock inoculated group samples showed evidence of DWV infection.

**Figure 4.**
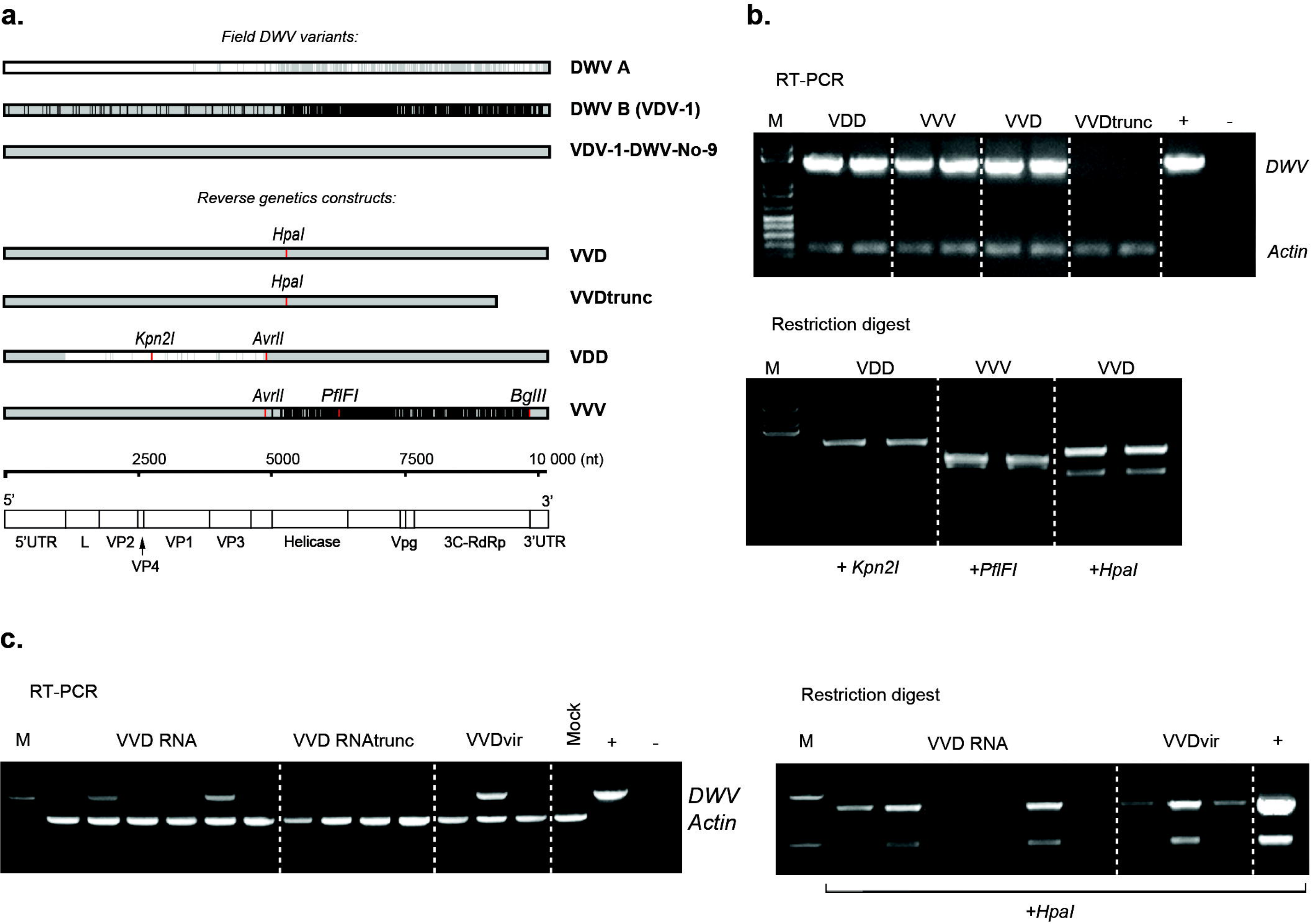
Morbidity of DWV in bumble bees. **A**. RT-qPCR analysis of DWV level in developed bumble bees injected with 10^6^ GE of DWV at white-eyed pupa stage. Individual values for each sample are shown with dots, lines represent mean ±SD. **B**. Percentage of visually normal (“*norm”*), discolored (“*disc”)* and non-viable (“*nv”)* bumble bees developed from pupae in VVD-injected (n=43) and PBS-injected (“Mock”, n=37). **C**. Phenotype of bumble bees developed from pupae in the incubator.

## Discussion

The term “pathogen spillover” describes the transmission of a pathogen from a reservoir host to a different species in a shared environment [44]. To achieve this, several conditions must be fulfilled; the reservoir host must be sufficiently abundant to guarantee exposure, pathogen prevalence must be high enough to ensure direct or environmental transmission, and the recipient host must be susceptible to infection via direct or indirect transmission [45].

Managed honey bee colonies are near-ubiquitous in many environments, including both rural and urban locations. On agricultural crops that require commercial pollination very high colony densities are achieved through migratory beekeeping. The combination of colony movements following the introduction of *Varroa destructor* resulted in the near-global distribution of the ectoparasite, with the concomitant spread of a range of honey bee viruses that are detrimental to colony survival [29]. Most important of these viruses is DWV which likely accounts for the majority of overwintering colony losses [46, 47]. DWV causes overt and lethal developmental defects after pupal inoculation and reduces the longevity of bees that do successfully emerge. Colonies that collapse due to high *Varroa*/DWV levels may be robbed-out by other insects including bumble bees, ants and wasps.

In addition to the cocktail of *Varroa* and DWV, honey bees are increasingly subject to stresses through a combination of limited dietary variation, repeated transportation to new pollination sites and exposure to agrochemicals, all of which are associated with increased susceptibility to pathogens and potential colony failure [48]. As a consequence, honey bees readily fulfill two of the requirements needed for pathogen spillover *i*.*e*. abundance in the environment and pathogen prevalence. In addition, due to the extensive foraging range of honey bees and the excretion of viable DWV in faeces [6], a high density of hives ensures widespread environmental contamination.

Notwithstanding the likely exposure of other species to the pathogen-laden environment, spillover also requires susceptibility of the species exposed and pathogen infectivity via a relevant transmission route. We exploited commercially available colonies of *Bombus terrestris* to determine if and how they could support infection by DWV. The buff-tailed bumble bee is a relevant model system in which to explore pathogen spillover; it is naturally found in the same environment as honey bees [4], and there are reports of DWV infection of wild-caught *B. terrestris* and several related species [4, 5, 11, 12, 22–24].

We developed a RG system for the two prototype strains of DWV (type A and B) and a hybrid previously reported to predominate in *Varroa*-infested colonies [32]. The synthesized virus genomes contain silent restriction sites that unambiguously distinguish the three types and – since they are unique to the RG cDNAs – allow discrimination from endogenous DWV. Direct inoculation of honey bee pupae with *in vitro* generated RNA resulted in DWV replication (Figure 2). Virus purified from these pupae caused symptomatic infection when reinoculated into honey bee pupae and accumulated to 10^10^ GE/μg of RNA (Figure S4), a level similar to that observed in *Varroa*-exposed pupae [25].

Using a similar strategy we demonstrated that bumble bee pupae could be infected when inoculated with *in vitro* generated DWV RNA. The resulting virus was purified, shown to be infectious for naïve bumble bee pupae when re-injected and deep sequencing of the virus population indicated no significant sequence changes from the originating cDNA (Figure S2). Infected pupae reached up to ∼10^9^ GE of DWV/μg of RNA two days post-injection (Figure 2A) and accumulation of the negative strand DWV RNA was confirmed by strand-specific qPCR (Figure S3). Therefore DWV is infectious for both honey bee and bumble bee pupae and adaptive changes are unlikely to create a bottleneck in any potential transmission between the species.

Direct recovery and amplification of DWV in bumble bees offers advantages to virologists and entomologists attempting to determine the significance of the limited differences between the reported strains. Since essentially all honey bee pupae are infected with DWV, perhaps with the exclusion of those from Australia [49], it is impossible to obtain truly clonal virus preparations. By recovering virus after RNA inoculation in bumble bees pure populations of DWV strains can be prepared for subsequent studies of virus pathogenesis and evolution.

Since *Varroa* does not parasitise bumble bee pupae it is difficult to rationalise direct injection as a potential transmission route. Therefore, considering robbing by bumble bees of collapsing honey bee colonies and the likely widespread DWV contamination in environments with large numbers of honey bees, we reasoned that oral transmission was a more likely route for virus acquisition. We therefore investigated oral susceptibility of larvae and adult bumble bees, fed directly or by extended exposure of bumble bee nests to virus-supplemented diet.

Direct feeding with 10^8^ GE of DWV enabled the detection of the negative strand of DWV, absent from the input virus preparation and indicative of virus replication, in ∼50% of larvae (Figure 2). In contrast, DWV fed adult bumble bees, or larvae and pupae harvested from nests supplemented with diet containing 10^8^ GE of DWV per bee, failed to provide any evidence for virus acquisition and transmission *per os*. Therefore, whilst larvae exhibit some susceptibility to infection by orally acquired DWV, it is clear that the threshold for infection may be high and that it cannot be achieved by extended feeding by adult bees throughout larval development.

Although we found no evidence for oral infection of adult bumble bees with DWV, we were able to demonstrate the presence of DWV in adult *Bombus* after direct virus injections. Adult bumble bees are able to support replication of DWV though, as with pupal inoculation with RNA, there may be a threshold (exceeded at 10^8^ GE) needed to achieve 100% infection.

In this study we allowed pupae to complete development and scored them phenotypically upon eclosion. Bumble bee pupae are susceptible to handling and survival rates (∼50%) were similar in virus- or mock-inoculated samples (Figure 4). Amongst the three phenotypically-distinct groups there was no difference in viral load in virus inoculated samples. Strikingly, the same three groups and the proportion of each were present in the mock-inoculated samples. No wing deformities were detected in eclosed bumble bees from either group.

It has previously been reported that DWV-positive field isolates of both *B. terrestris* and *B. pascuorum* have been identified with wing deformities characteristic of those seen in DWV-infected honey bees [7]. However, these studies did not demonstrate DWV replication or reproduce symptoms by DWV inoculation. In honey bees apart from DWV infection, wing malformation can be caused during development by injury, hormonal disorders, intoxication or absence of cocoon [37]. While laboratory inoculations represent the gold standard for virus infectivity assays this approach probably does not recapitulate an environmentally meaningful infection route. Compared to their independent effect a combination of stressors is suggested to introduce a greater threat to wild pollinators in their native environment [50]. For example, exposure to clothianidin - a neonicotinoid insecticide - was found to have a negative impact on honey bee immune status and promote DWV infection [51]. In *Bombus terrestris* condition mediated virulence of Slow bee paralysis virus upon starvation has been demonstrated [52]. This study uses a reverse genetic approach to investigate pathogen spillover from honey bees to bumble bees. Whilst clear evidence is obtained to confirm DWV replication in bumble bee larvae, pupae and adults, we were unable to demonstrate a compelling route by which transmission would likely occur in the natural environment. The levels of virus required to orally infect bumble bee larvae are significantly higher than have been reported in environmental pollen samples [53]. Indeed, the levels required are likely higher than larvae are ever exposed to. In contrast, adult bumble bees may experience very high virus levels in collapsing honey bee colonies while robbing. However, adult bumble bees feeding on virus-supplemented syrup remain uninfected and – importantly – are unable to transmit infectious virus to their developing larvae. Further studies will be required to determine whether the gut environment of adult bumble bees is sufficiently hostile to DWV that the virus is inactivated *e*.*g* by gut proteases, or if there are other factors that restrict infection and replication of DWV in bumble bees.

Previous reports strongly underline the haplotype identity of DWV between honey bee and bumble bee populations [4, 24]. Published data show that majority of the virus is located in the gut, which can be interpreted either as a restriction of infection to the digestive tract or as presence of DWV in the ingested material [22]. The results obtained from this study provides a strong indication that oral acquisition of virus from a contaminated environment does not represent an effective DWV transmission route between *Apis* and *Bombus*. Bumble bees are known to carry their own mite parasites, with *Apicystis bombi* and *Crithidia bombi* found in honey bee collected pollen [54]. However, these mites do not parasitise honey bees and lack feeding behaviour similar to Varroa or the capacity for virus vectoring. Therefore, a route for productive DWV spillover to bumble bees from infected honey bees remains to be determined. We propose that ubiquitous detection of DWV RNA in geographically related bumble bees primarily reflects the carryover material originating from virus prevalence in the abundant honey bee population, whereas incidence of pathogen spillover is limited to the cases of conditioned infection. Further studies on defining the stressors causing DWV infectivity in bumble bees are required in order to estimate the actual impact of DWV on this important group of pollinators.

## Materials and Methods

### Honey bees

All honey bee (*Apis mellifera*) brood were collected from the University of St Andrews research apiary. All hives used for sampling were routinely treated for *Varroa* with an appropriate miticide. Pupae were extracted from the comb and maintained in the incubator set at 34.5°C with 90% relative humidity.

### Bumble bees

Bumble bees *(Bombus terrestris audax*; Biobest, Belgium) were maintained in the laboratory in isolated nests supplemented with feeders containing 50% aqueous sucrose solution. Nests were regularly fed with bee pollen from a DWV-free region (Saxonbee, Australia). Separately treated adult bumble bees were maintained in cages at RT and fed *ad libidum* on pollen/syrup. Pupae and larvae were extracted from the brood cells and transferred into the incubator set at 30.5°C with 90% relative humidity. Larvae were fed a diet consisting of 25% (v:v) ground pollen and 25% sucrose solution (w:v) in H_2_O.

### RG system and *in vitro* RNA synthesis

All RG constructs in this study were based on a cDNA clone of a recombinant DWV variant VDV-1_VVD_ (GenBank HM067438.1). The initial pVVD construct was made using a similar protocol described for previously published reverse genetics systems of DWV [37, 39]. Briefly, the full-length cDNA was cloned in a pBR-derived plasmid vector, flanked at the 5’ end by a T7 promoter and hammerhead ribozyme [55], and at the 3’ end by a *Pme* I restriction site following the poly-A tail. VDD (DWV-A) and VVV (DWV-B) constructs were built by substituting parts of pVVD, with the DWV-A capsid-coding and DWV-B non-structural protein-coding cDNA sequences based on published data [27, 40, 41] and produced by custom gene synthesis (IDT, Leuven, Belgium). Unique synonymous restriction sites were introduced into each cDNA sequence. All resulting plasmids were verified by Sanger sequencing.

RNA was generated *in vitro* from linearised templates, using the T7 RiboMAX™ Express Large Scale RNA Production System (Promega). Full length and truncated templates were linearised with *Pme* I or *Nru* I (nt 9231) respectively. Truncated RNA transcripts were polyadenylated using the poly(A) Tailing Kit (Thermo Fisher Scientific) to ensure their stability. After purification, RNA transcripts were eluted in RNAse free H_2_O, their integrity confirmed by gel electrophoresis, and stored at −80°C.

### RNA and virus injections

RNA and virus injections of 5 and 10 μl were performed with insulin syringes (BD Micro Fine Plus, 1 ml, 30 G) into honey bee and bumble bee pupae respectively. Up to 10 μg (equivalent to 1.7×10^12^ of DWV RNA copies) of *in vitro* transcribed RNA was injected individually into pupae. Truncated VVD transcript injections were used as a negative control. All RNA-injected pupae were analyzed at 72 h post injection. Injections of pupae or low temperature-immobilized adult bees with virus stocks diluted in PBS were performed using the same technique as for RNA transcripts.

### Virus stocks

DWV stocks were prepared from pupae injected with *in vitro* transcribed RNA. Homogenized tissue was diluted with sterile PBS in 1:1 (w:v) ratio and centrifuged at 13 000 g, 4°C for 10 min. Supernatant was filter sterilized with 0.22 μM filter (PES, Millipore) and treated with RNase A to destroy all unencapsidated RNA. RNA was extracted from 100 μl of the virus stock using RNeasy kit (Qiagen) and analyzed by RT-qPCR. Apart from the virus inoculum used for testing the infectivity of the *Bombus*-derived DWV, all virus stocks were prepared from honey bee pupae.

### Oral inoculation and bumble bee nest feeding

Individual bumble bee larvae were placed into 96 well plates containing 2 μl of pollen/syrup mixed with DWV inoculum. DWV feeding was delivered once on the first day of the experiment. Fresh diet without virus was added individually after virus-containing food had been consumed. On the fifth day larvae were snap-frozen in liquid nitrogen and stored at −80°C.

Adult bumble bees were fed with virus in groups of 20 on the first and on the second day of the experiment. Total amount of DWV supplied with each of the two virus feedings was 2.2×10^9^ and 2.2×10^8^ DWV GE for 10^8^ and 10^7^ groups respectively. Bumble bees were snap-frozen in liquid nitrogen and analyzed for the presence of DWV RNA 7 days after DWV feeding.

For the colony-scale inoculations three bumble bee nests each containing a mated queen and 120-125 adult bees were used. Prior to the start of the experiment all brood was removed from the nests. Virus feeding was delivered daily by replacing the nest feeder with a tube containing 2 ml of sucrose solution supplemented with DWV, previously confirmed to be infectious for bumble bee pupae by direct injection. Each nest received a daily dose of the virus corresponding to 2.4×10^10^ DWV GE. The virus-containing solution was fully consumed (<3-4 h) before replacement with the regular sucrose feeder. DWV feeding continued for 4-6 weeks and stopped when the first group of larvae developed during the virus-feeding period approached pupation. Upon termination of the experiment all brood (eggs, larvae and pupae) and 10 adult workers were sampled from each nest and tested for the presence of DWV.

### RNA extraction, reverse transcription and PCR

Samples were homogenized with a Precellys Evolution homogenizer (Bertin Instruments, France). Total RNA was extracted using the GeneJet RNA Purification Kit (Thermo Scientific). cDNA was prepared with qScript cDNA Synthesis Kit (Quanta Biosciences) from 1 μg of total RNA following the manufacturer’s protocol. Detection of DWV and actin mRNA, used as an internal RNA quality control, was carried out by end-point PCR with *Taq* DNA polymerase (New England Biolabs) and 2 μl of cDNA. DWV_RTPCR primers were designed to detect all three DWV variants under study (Table S1). To amplify the regions containing restriction site tags in VDD and VVV Kpn2I_F/R and PflFI_F/R primer pairs were used respectively. PCR cycling conditions were 30 cycles of 95°C (15 seconds), 55°C (15 seconds), 68°C (2 minutes) with an initial 95°C step (30 seconds) and a final extension at 68°C (5 minutes). PCR samples were analyzed on a 1% agarose gel and when required, PCR products were subjected to restriction digest prior to loading on the gel.

The quantification of DWV genome copies was performed by SYBR-Green Real-Time Quantitative PCR (qPCR) using Luna Universal qPCR master mix (New England Biolabs), 0.25 μM forward and reverse DWV_qPCR primers and 2 μl of cDNA. The following thermal profile was used: 1 min at 95°C, followed by 40 cycles of 15 seconds at 95°C and 30 seconds at 60°C with a final post-amplification melting curve analysis step. DWV titres were calculated by comparison of the resulting Ct value to the standard curve generated from a serial dilution of the VVD cDNA qPCR standard prepared by reverse transcribing the RNA transcript.

### Negative strand assay

Strand-specific detection of DWV RNA was performed as described earlier [25]. Briefly, 1 µg of total RNA was used in reverse transcription reaction carried out with Superscript III reverse transcriptase (Invitrogen) and the adapter extended primer designed to anneal to the negative strand RNA of DWV. The PCR step was carried out by *Taq* DNA polymerase (NEB) using forward primer identical to the adapter sequence (Table S1) and DWV_RTPCR_R primer. PCR was run for 35 cycles in the same conditions as described above.

## Acknowledgements

We would like to express our gratitude to Robert J Paxton for providing a reference sequence for the construction of DWV-B RG system. This work was supported by grant funding from BBSRC BB/M00337X/2 and BB/I000828/1.

