## Supplementary information for "Deformed Wing Virus spillover from honey bees to bumble bees: a reverse genetic study" (from another version of this preprint: version 3, 2020-06-19)

### Evidence for and against Deformed Wing Virus spillover from honey bees to bumble bees: a reverse genetic analysis.

**Supplementary materials**

**Table S1. Primers used in this study.**

| Name | Sequence (5'-3') | cDNA binding site | PCR product size | Reference |
|---|---|---|---|---|
| **DWV_RTPCR_F** | ATTGATCATTGTATGTTTACCTTC CCTTG | 4021-4049 | 1606 bp | [46] |
| **DWV_RTPCR_R** | GCACGTAAGAGCTCGCTGCATA | 5606-5627 | | [46] |
| **DWV_qPCR_F** | ATATCACTTGGCGACGCAAC | 4950-4969 | 176 bp | [46] |
| **DWV_qPCR_R** | CCAATCTTTAAATTGTTTCGGTTT TTGAGC | 5098-5126 | | [46] |
| **Kpn2I_F** | ATCCAGTGCAGGCAAAACCA | 1763-1782 | 1545 bp | [46] |
| **Kpn2I_R** | TACCGCCCCGTACGTACACA | 3288-3307 | | [46] |
| **PflFI_F** | TAATCCTGCAGCGCGACTCT | 5247-5266 | 1530 bp | [46] |
| **PflFI_R** | GAACGCTCGTGGACATACGC | 6757-6776 | | [46] |
| **DWV(-RNA)_RT** | CTTGGTTAGCTGTGTTGCAGTTGC TGTAGTTAAGCGGTTATTAGAA | 4925-4947 | 120bp | [25] |
| **DWV(-RNA)_RT R** | CTGTAGTTAAGCGGTTATTAGAA | 4925-4947 | | [25] |
| **388 (adapter)** | CTTGGTTAGCTGTGTTGCAGTTG | | | [25] |
| ***Apis mellifera* actin F** | AGGAATGGAAGCTTGCGGTA | 919–938 | 181 bp | [32] |
| ***Apis mellifera* actin R** | AATTTTCATGGTGGATGGTGC | 1099–1079 | | [32] |
| ***Bombus terrestris* actin F** | AGGGTGTGATGGTCGGTATGG | 261-281 | 951 bp | *This study* |
| ***Bombus terrestris* actin R** | GAGATCCACATCTGTTGGAAGG | 1191-1212 | | *This study* |
| **DWV FG RP1** | TACGCGAGTAACACCTAAC | 10068-10086 | 10068bp | *This study* |
| **DWV FG FP4** | GCGAATTACGGTGCAACTAAC | 19-39 | | *This study* |

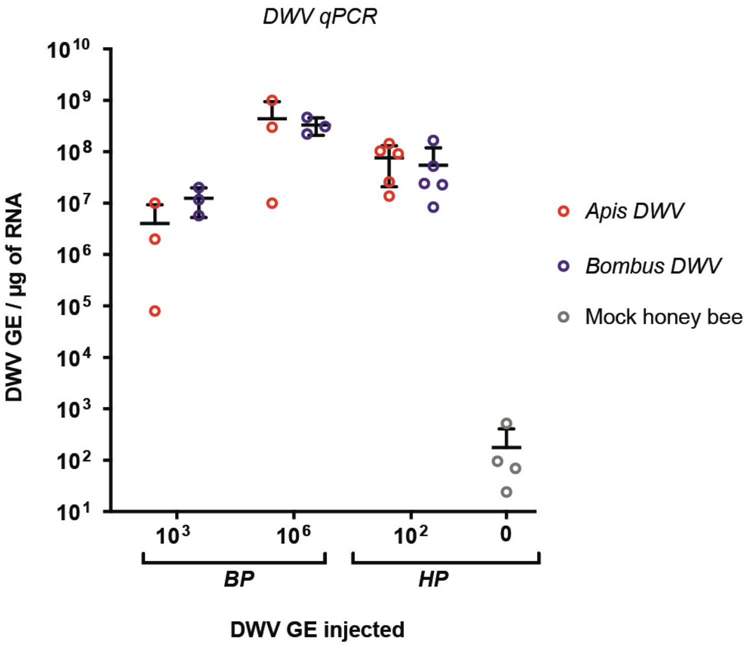


**Figure S1. Infectivity of *Bombus* and *Apis*-derived DWV.** qPCR analysis of DWV accumulation in bumble bee (BP) and honey bee (HP) pupae injected with VVD DWV inoculum prepared from pupae transfected with *in vitro* synthesized RNA. "*Apis*" and "*Bombus*" DWV stocks were prepared from honey bee and bumble bee pupae tissues respectively. Each value corresponds to an individual sample analyzed, error bars show mean ±SD. After injection bees were incubated for 48 and 24 h before analysis for bumble bee and honey bee pupae respectively. GE - genome equivalents.

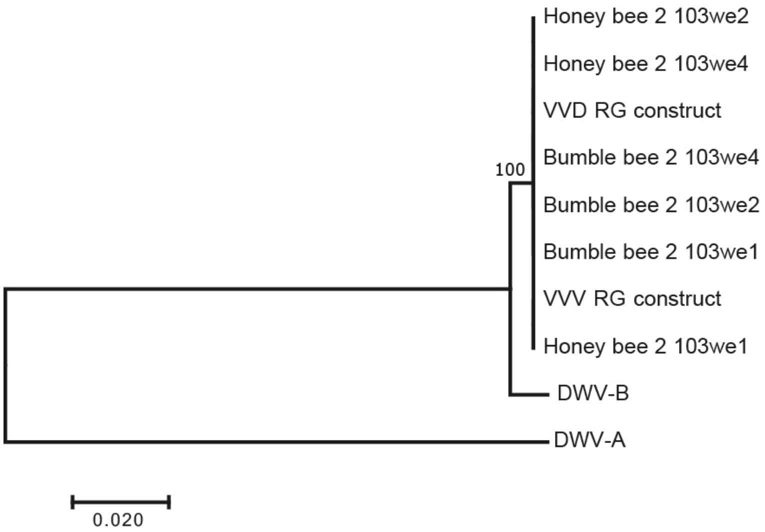


**Figure S2. Phylogenetic analysis of DWV genomic RNA in first generation virus samples from injected honey bees and second generation virus passaged in injected bumble bees**. DWV RNA isolated from pupae injected with virus at white-eyed stage (P0-P1 stage for bumble bees) was sequenced using Illumina and subsequently analysed by ShoRAH to identify any adaptive sequence changes which may have occurred after virus passaging in bumble bees. First generation DWV sequences were obtained from honey bee pupae inoculated with virus stock prepared from RG RNA-injected honey bees, second generation DWV sequences were obtained from bumble bee pupae inoculated with DWV stock prepared by an additional round of passaging of the virus in bumble bee pupae (re-injection of the virus stock obtained from RNA-injected bumble bee samples). The phylogenetic tree shows that the top matches from ShoRAH all align identically to the originally injected variant (VVD RG construct), indicating no adaptive change has taken place in the non-Apis host. Reference sequences for DWV type A and type B are included. A 500 bootstrap neighbour-joining tree was generated.

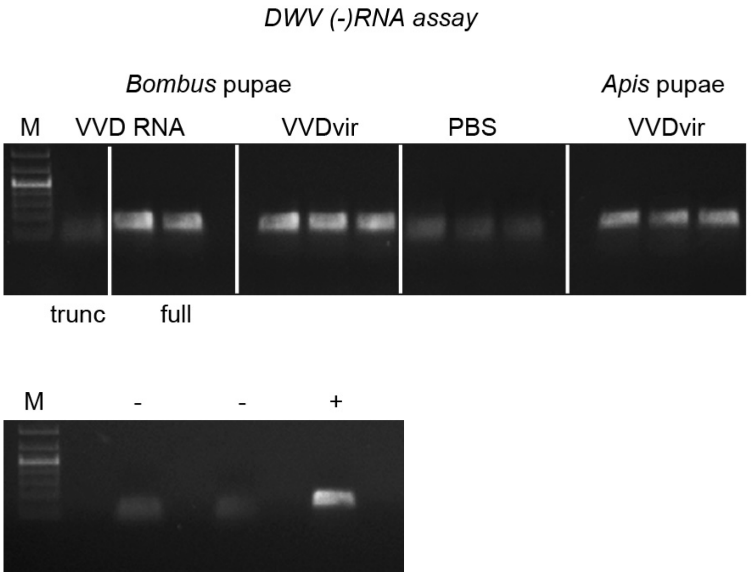


**Figure S3. Detection of the DWV (-)RNA strand in bumble bee and honey bee pupae**. Strand-specific reverse transcription and PCR assay for (-)RNA of DWV in pupae injected with full-length ("full") and truncated ("trunc") VVD RNA, VVD virus inoculum ("VVDvir") or PBS (negative control). RNA-injected pupae were analyzed 3 days post-injection. Virus-injected pupae were analyzed 48 and 24 h post-injection for bumble bees and honey bees respectively. "+" and "-" - positive and negative PCR controls, "M" - molecular weight DNA marker. RG origin of the PCR products for all DWV-positive samples was confirmed by restriction enzyme digest.
